# Methane-fed microbial communities enriched from field-grown rice support diverse heterotrophic bacteria

**DOI:** 10.1101/2025.06.09.658666

**Authors:** Elena K. Perry, Aqib Hasnain, Benjamin J. Cole, Hans K. Carlson, Adam M. Deutschbauer, Dawn Chiniquy

**Affiliations:** Environmental Genomics and Systems Biology Division, Lawrence Berkeley National Laboratory, Berkeley, CA, USA; DOE Joint Genome Institute, Lawrence Berkeley National Laboratory, Berkeley, CA, USA

## Abstract

Rice paddies naturally host aerobic methanotrophic bacteria, due to the production of methane in flooded soils. However, relatively little is known about how the activity of methanotrophs impacts the structure of the broader microbial community in this globally important agricultural environment. To address this question, we passaged 51 aerobic microbial enrichment cultures from rice rhizosphere, root, and stem samples in a chemically-defined medium with methane as the primary carbon source and electron donor. We profiled the cultures over time by 16S rRNA gene amplicon sequencing and sequenced the genomes of 40 isolates from the enrichments to gain functional insights. Taxa whose relative abundance increased during community growth on methane represented more than a dozen families, many of which are not known to utilize methane or other one-carbon substrates. Despite the selective pressure imposed by the culture condition, the final community structures were taxonomically varied rather than converging to a common composition. Genomic analysis of the sequenced isolates revealed considerable variation in likely carbon source utilization repertoires, as well as the capacity for nitrogen fixation or denitrification. Taken together, these findings support the view that methanotrophy represents a key link in the microbial food web of rice fields, with the potential for downstream effects on the abundance and activity of a wide range of community members.

**Importance:** Rice paddies produce the primary staple food for more than half of the world’s population, but are also a major source of methane emissions. Methane-oxidizing bacteria known as methanotrophs naturally occur in rice paddies, where they consume methane that would otherwise escape to the atmosphere. Enhancing the activity of native methanotrophs could improve the sustainability of rice cultivation. However, large gaps remain in our knowledge of how increased methanotrophic activity could impact other members of rice paddy microbial communities, leaving open the possibility of unintended effects on other important microbial ecosystem functions, such as nitrogen cycling. This study sheds light on the taxonomic range and metabolic capacities of rice-associated bacteria that can benefit from the activity of native methanotrophs, laying the foundation for a broader understanding of how methanotrophs impact microbiome assembly and nutrient cycling in rice paddies.

## Introduction

Microbes that oxidize one-carbon compounds, such as methane and methanol, play a key role in global nutrient cycles. In particular, aerobic methanotrophs in the phylum Pseudomonadota are the most widespread functional guild contributing to microbial methane oxidation. Aerobic methanotrophs are classically divided into two major groups, type I (Gammaproteobacteria) and type II (Alphaproteobacteria); type I methanotrophs use the ribulose monophosphate (RuMP) pathway for carbon assimilation into biomass, while type II use the serine pathway. Because these bacteria can use methane as a sole carbon and energy source, they are found in a variety of environments where methane is abundant, such as wetlands, freshwater lake sediments, and landfill cover soils (1). Besides contributing to carbon cycling via methane oxidation, methanotrophs may also play an important role in the global nitrogen cycle: many proteobacterial methanotrophs are capable of nitrogen fixation (2), although conversely, methanotrophs have also been shown to stimulate the activity of denitrifying bacteria (3, 4).

Consistent with the latter observation, it is thought that methanotrophs often interact metabolically with other members of their native microbial communities (5). Methanotrophs are notoriously difficult to isolate (6), and culturing methanotrophs in the presence of other heterotrophic bacteria frequently increases methane oxidation rates and overall biomass yields (7, 8). Such partnerships may benefit the methanotrophs by preventing the accumulation of inhibitory metabolic byproducts or providing essential vitamins (7, 9). Methanotrophs can in turn provide benefits to their neighbors. Studies using stable-isotope probing (SIP) have demonstrated that carbon derived from methane by methanotrophs in a variety of environments can end up in the biomass of diverse other organisms (10–15). Thus, methanotrophs contribute to local microbial food webs by transforming methane into other, more bioavailable carbon sources.

While the existence of metabolic exchanges between methanotrophs and other bacteria is well-established, relatively little is known about how methanotrophs influence the microbial communities of methane-rich wetland environments, including rice paddies (16). Recently, methanotrophs have been shown to stimulate both nitrogen fixation and denitrification in rice paddies, depending on the environmental conditions (12, 17), but the full range of their impact on the rice microbiome remains unclear. Given the roles of methanotrophs in both carbon and nitrogen cycling, an improved understanding of their ecological influence in rice paddies could have significant implications for improving the productivity and sustainability of rice cultivation. This issue is of global importance since rice is one of the world’s most widely-consumed staple crops (18). The most concrete evidence that specific taxa might preferentially benefit from methane-derived carbon in rice fields has come from DNA-SIP studies on microcosms of paddy soils or rice roots (10, 12). In these studies, the bacteria that incorporated the ^13^C label mainly belonged to the Alphaproteobacteria and Betaproteobacteria, although the results varied at lower taxonomic levels. However, the sensitivity of DNA-SIP is limited and the results can be significantly affected by experimental parameters such as incubation time (19), thus motivating the use of complementary approaches to determine the range of microbial taxa that may benefit from methanotrophic activity in rice fields.

Here, we performed 16S rRNA gene amplicon-based profiling on methane-fed enrichment cultures, complemented by whole-genome sequencing of strains isolated from the enrichments, to gain new insights into how methanotrophs from rice paddies influence their native communities. Specifically, we addressed the following questions: 1) What is the taxonomic diversity of rice-associated bacteria that can be supported by methane-derived carbon? 2) To what extent is the community composition of methane-fed enrichment cultures affected by external variables such as the part of the rice plant used for the inoculum? 3) Is the impact of methanotrophs on the rice microbiome predictable, as indicated by recurring community assembly patterns (if any) during growth on methane as the sole exogenous carbon source? 4) What carbon sources are the most likely drivers of non-methanotrophic growth in methane-fed, methanotroph-anchored communities?

## Results

### Methane-fed enrichment cultures maintained high alpha diversity

We established 51 enrichment cultures in chemically-defined media with methane as the sole electron donor and oxygen as the major electron acceptor, inoculated with rice rhizosphere, root, and stem samples collected from two field locations in California, as described in Materials and Methods. The cultures were sampled weekly over 12 weeks for 16S rRNA gene amplicon-based profiling of community composition, resulting in a total of 534 sequenced samples (excluding those that failed to meet our threshold for minimum reads; see Materials and Methods); the complete amplicon sequence variant (ASV) abundance table and sample metadata are available in Tables S1 and S2, respectively. At the final time point, 16 families belonging to three phyla (Pseudomonadota, Bacteroidota, and Verrucomicrobiota) achieved a relative abundance of >10% in at least two enrichments (Fig. 1A). Two additional higher-level taxonomic groups, consisting of ASVs that could not be classified beyond the order Rhizobiales or the domain Bacteria, met the same criteria for abundance and prevalence. By far the most prominent families across the final enrichment communities, when considering both prevalence and relative abundance, were Comamonadaceae (>10% in 33/51 enrichments, mean relative abundance 16.3%) and Methylocystaceae (>10% in 29/51 enrichments, mean relative abundance 22.4%). Comamonadaceae does not contain any known methanotrophs, but Methylocystaceae represents the Type II methanotrophs. Type I methanotrophs, represented by the family Methylococcaceae, were less prevalent in the final enrichments (>10% in 4/51 enrichments).

**Figure 1.**
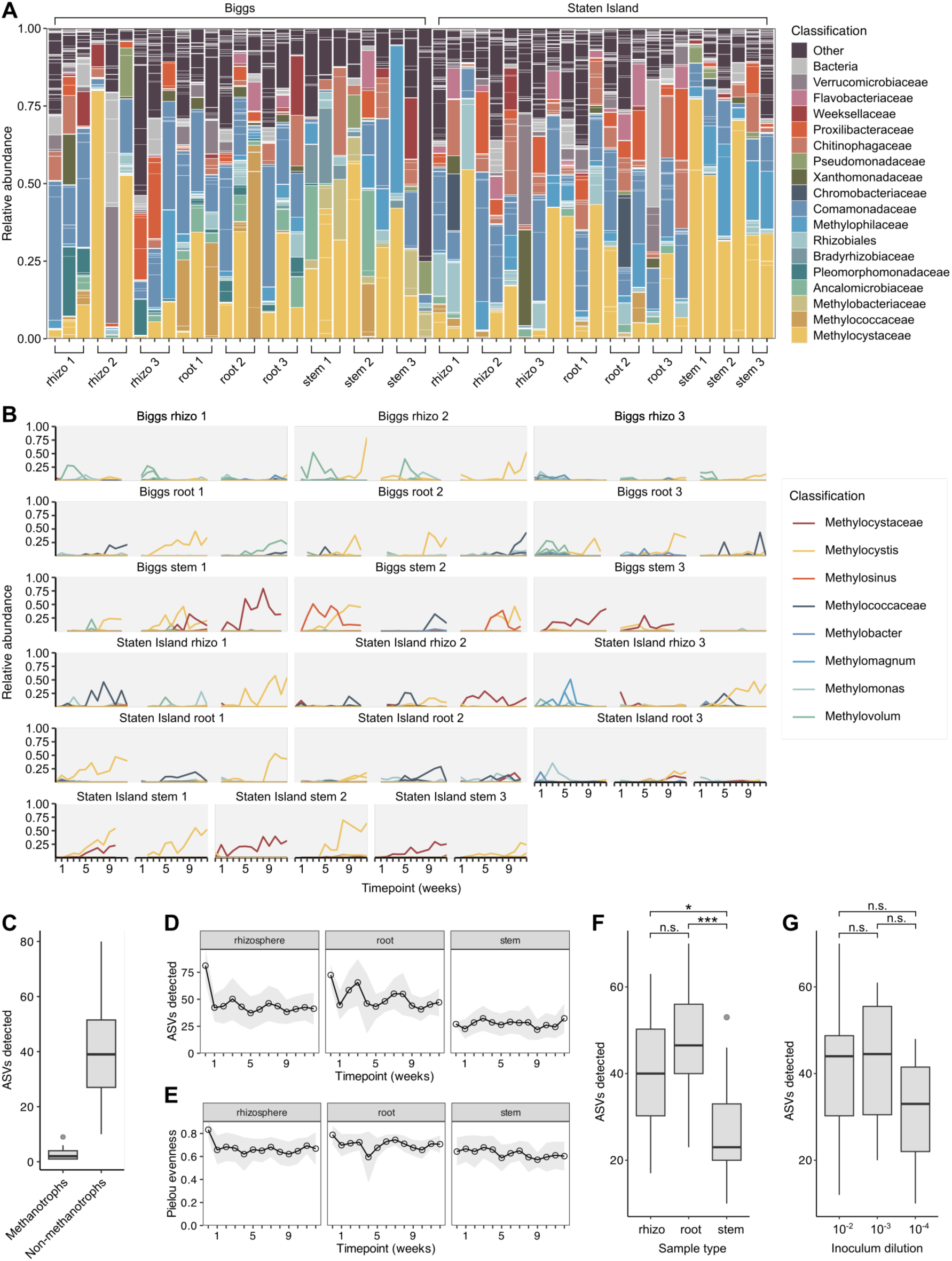
Taxonomic diversity and dynamics of methane-fed enrichment cultures. **A)** Taxa bar plot showing the overall diversity and major families in the enrichment cultures at the final sequenced timepoint. Individual ASVs within each family are separated by white lines. **B)** Relative abundance of putative methanotroph ASVs over time in each enrichment. Under each header (e.g. “Biggs rhizo 1”), the enrichments are arranged from left to right in order of increasing inoculum dilution (i.e., 10^−2^, 10^−3^, 10^−4^). **C)** The number of detected ASVs that were classified as methanotrophs or non-methanotrophs in each enrichment at the final timepoint. **D-E)** The number of ASVs detected (D) or Pielou evenness (E) of the enrichments over time, separated by sample type. **F)** The number of ASVs detected in the enrichments at the final timepoint, separated by sample type or by inoculum dilution. * *p* < 0.05, *** *p* < 0.001, ANOVA followed by Tukey HSD.

Interestingly, while all of the final enrichment communities contained a detectable level of at least one putative methanotroph ASV classified as Methylocystaceae or Methylococcaeae, methanotrophs were not always the most abundant community members. Methanotrophs represented the majority of reads in just eight out of the 51 final communities. In 16 out of the 51 enrichments (nearly 1/3) the final total methanotroph relative abundance was <10%, and the lowest final total methanotroph relative abundance was just 0.2%. Inspecting the relative abundances of putative methanotroph ASVs over time revealed that in many of the enrichments, methanotroph abundances fluctuated or dropped after reaching a peak prior to the final time point, rather than steadily increasing over time—especially for many of the Type I methanotrophs (Fig. 1B). Additionally, in most of the enrichments only one or two methanotroph ASVs were abundant at any given time (Fig. 1B-C). By contrast, individual enrichments supported up to 80 non-methanotroph ASVs (with a median of 39) at the final time point (Fig. 1C).

We also assessed how overall alpha diversity was affected by time, sample type, and initial inoculum dilution. For rhizosphere and root enrichments, the number of ASVs observed per enrichment initially dropped noticeably compared to the inoculum, but subsequently fluctuated around the value reached by the end of week one, rather than continuing to decline, despite the 1:10 dilution transfers that were performed after week four and week eight (Fig. 1D). For stem enrichments, the number of observed ASVs dropped only slightly in the first week, and fluctuated around the original value thereafter (Fig. 1D). Community evenness followed generally similar patterns over time for the different sample types (Fig. 1E). At the final time point, the number of observed ASVs was significantly lower for stem enrichments compared to rhizosphere and root enrichments, but did not differ between the latter two groups (Fig. 1F). Initial inoculum dilution had no statistically significant effect on the final number of observed ASVs (Fig. 1G).

### Beta diversity arose primarily from individual variation between enrichments rather than external variables

The taxa bar plots for the final time point indicated considerable variability in community composition across the enrichments (Fig. 1A). We therefore performed additional analyses to assess the impact of three external variables on beta diversity: time since inoculation, field site of origin, and original sample type. PERMANOVA analysis indicated that time had a significant effect on beta diversity regardless of whether ASV abundances and phylogenetic relationships were accounted for (*p* = 0.001 for both Jaccard and weighted UniFrac distances) (Table 1). However, the effect was relatively small (R^2^ = 0.04 and 0.08 for Jaccard and weighted UniFrac distances, respectively), and the respective PCoA plots suggested that the effect was not particularly consistent over time or across different enrichments (Fig. 2A). Indeed, the time to reach a stable community composition varied across the enrichments, with some appearing to largely stabilize following the first 1:10 dilution transfer after week four; some experiencing another dramatic shift following the second 1:10 dilution transfer after week eight; and a few appearing to undergo a more or less continuous shift starting after the first transfer and continuing to the end of the experiment (Fig. S1). Additionally, PERMDISP analysis indicated that the sample dispersions differed significantly across different time points (*p* = 0.001 for both Jaccard and weighted UniFrac distances) (Table 1), which can cause a significant PERMANOVA result even if there is no difference in the group centroids.

**Figure 2.**
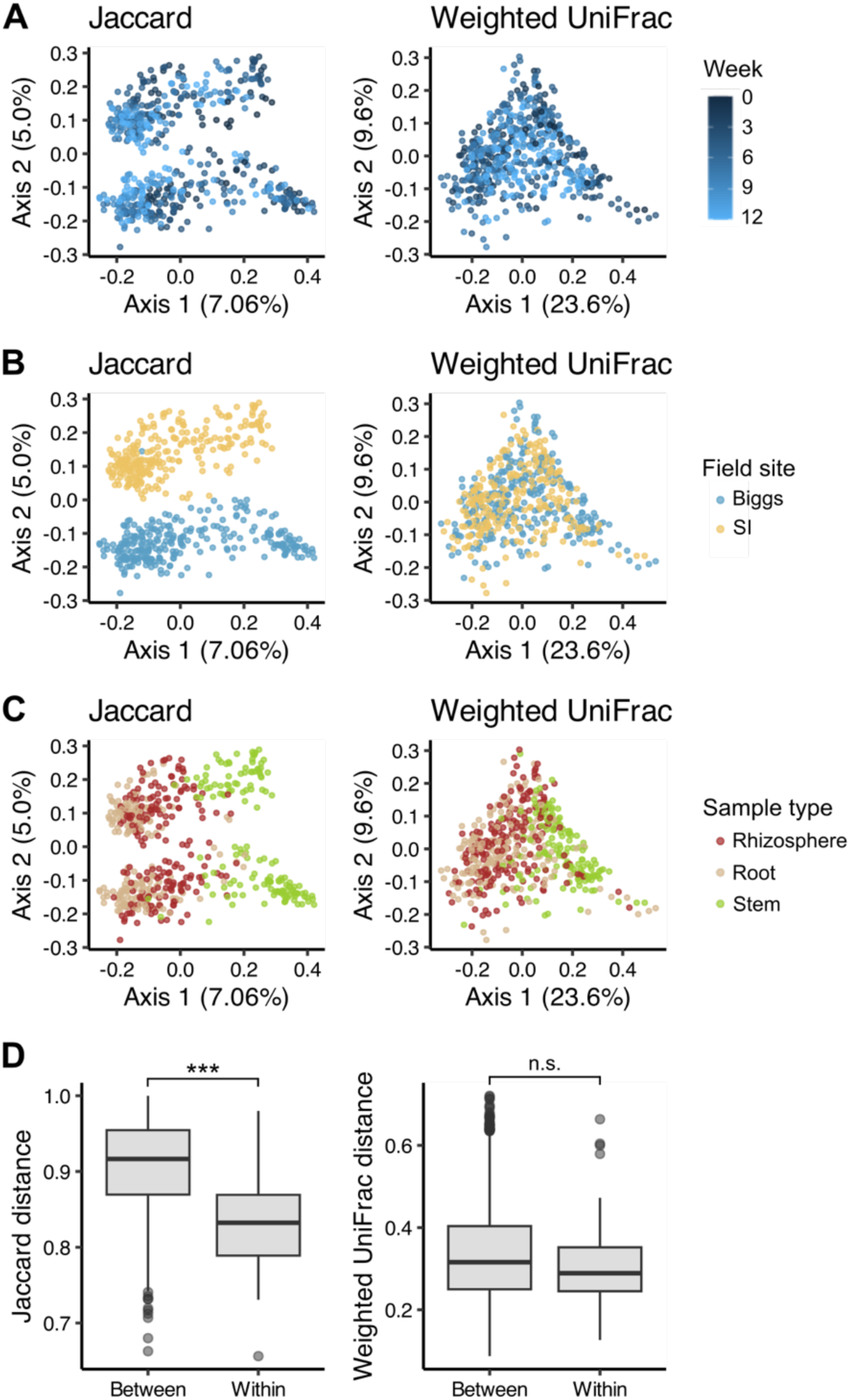
Impact of external variables on enrichment community beta diversity. **A)** PCoA plots of all sequenced enrichment samples, colored by time point. **B)** The same plots as in (A), but colored by field site. **C)** The same plots as in (A), but colored by sample type. **D)** Jaccard and weighted UniFrac distances for enrichments inoculated from different replicate plant parts (“between”) versus enrichments inoculated from the same replicate plant part (albeit at different dilutions; “within”).

**Table 1.** PERMANOVA and PERMDISP statistical test results.

| Distance metric | Test | Independent variable | R<sup>2</sup> | Pseudo-F | Pr(>F) |
| --- | --- | --- | --- | --- | --- |
| Jaccard | PERMANOVA | Timepoint | 0.04 | 2.0427 | 0.001 |
| Weighted UniFrac | PERMANOVA | Timepoint | 0.08 | 3.8124 | 0.001 |
| Jaccard | PERMDISP | Timepoint | NA | 5.8664 | 0.001 |
| Weighted UniFrac | PERMDISP | Timepoint | NA | 8.0389 | 0.001 |
| Jaccard | PERMANOVA | Field site | 0.05 | 25.623 | 0.001 |
| Weighted UniFrac | PERMANOVA | Field site | 0.03 | 14.035 | 0.001 |
| Jaccard | PERMDISP | Field site | NA | 50.335 | 0.001 |
| Weighted UniFrac | PERMDISP | Field site | NA | 19.947 | 0.001 |
| Jaccard | PERMANOVA | Sample type | 0.07 | 20.108 | 0.001 |
| Weighted UniFrac | PERMANOVA | Sample type | 0.13 | 39.414 | 0.001 |
| Jaccard | PERMDISP | Sample type | NA | 51.113 | 0.001 |
| Weighted UniFrac | PERMDISP | Sample type | NA | 17.582 | 0.001 |
| Jaccard | PERMANOVA | Enrichment ID | 0.52 | 10.412 | 0.001 |
| Weighted UniFrac | PERMANOVA | Enrichment ID | 0.44 | 7.4642 | 0.001 |
| Jaccard | PERMANOVA | Replicate (marginal effect*) | 0.01 | 7.9084 | 0.001 |
| Weighted UniFrac | PERMANOVA | Replicate (marginal effect*) | 0.01 | 7.9996 | 0.001 |
| Jaccard | PERMANOVA | Dilution (marginal effect*) | 0.02 | 6.1119 | 0.001 |
| Weighted UniFrac | PERMANOVA | Dilution (marginal effect*) | 0.02 | 7.8126 | 0.001 |

\*Marginal effect indicates the effect of the term after taking into account all other terms in the model (the terms were timepoint, field site, sample type, replicate, and dilution).

The field site of origin (Biggs versus Staten Island) also had a statistically significant effect on community composition (PERMANOVA *p* = 0.001 for both Jaccard and weighted UniFrac distances, permuting only within each time point given the significant effect of time) (Table 1). However, while the PCoA plot based on Jaccard distances revealed an almost perfect separation between enrichments from the two fields, this was not evident with weighted UniFrac distances (Fig. 2B), and the overall effect of field site was again quite small in both cases (R^2^ = 0.05 and 0.03 for Jaccard and weighted UniFrac distances, respectively). Original sample type (rhizosphere, root, or stem) also had a significant effect on community composition (PERMANOVA *p* = 0.001 for both Jaccard and weighted UniFrac distances, permuting only within each time point), with slightly larger effect sizes compared to field site (R^2^ = 0.07 and 0.13 for Jaccard and weighted UniFrac distances, respectively). Indeed, while the PCoA plots suggested a high degree of overlap among rhizosphere and root enrichments, the stem enrichments clustered separately to a moderate degree based on both Jaccard and weighted UniFrac distances (Fig. 2C). However, PERMDISP indicated that as with the effect of time, sample dispersions also differed for the two field sites and across the different sample types (*p* = 0.001 for both Jaccard and weighted UniFrac distances) (Table 1).

In contrast to the relatively small effects of the above variables, by far the greatest proportion of the beta diversity was explained by individual enrichment identity, i.e. the unique combination of field site, sample type, replicate, and dilution (PERMANOVA *p* = 0.001, R^2^ = 0.44 for weighted UniFrac) (Table 1). The marginal effects of replicate and dilution were very small (R^2^ = 0.01 and 0.02 respectively), although still statistically significant (*p* = 0.001) (Table 1). We therefore also asked whether enrichments derived from the same original sample but with different inoculum dilutions (“within sample”) ended up more similar to each other at the final time point than enrichments derived from different original samples (“between sample”). In the case of Jaccard distances (which take into account only the presence/absence of each ASV), there was a statistically significant difference for the “within sample” versus “between sample” comparison (Fig. 2D). This intuitively makes sense as enrichments derived from the same original sample would be expected to share a higher proportion of ASVs, given the lack of opportunities for dispersal between the enrichments over time. Notably, however, based on the weighted UniFrac metric, which provides an arguably better representation of overall differences in community structure, there was no statistically significant difference for “within sample” versus “between sample” distances.

### Stochastic processes dominated the enrichment community assembly

Given the high degree of variability in final community composition, even across individual enrichments derived from the same original sample, we hypothesized that stochastic processes had contributed significantly to community assembly in the enrichments. We therefore sought to quantitatively infer the contribution of stochastic versus deterministic ecological processes (i.e. drift versus selection), using two related statistical methods known as QPEN (“Quantifying assembly Processes based on Entire-community Null model analysis”) and iCAMP (“infer Community Assembly Mechanisms by Phylogenetic-bin-based null model analysis”) (20–22). Both approaches are founded on the observation that more closely related species tend to occupy more similar niches; thus, if the abundance-weighted phylogenetic relationships between two communities’ members are on average significantly closer than expected by chance given the full set of taxa observed across all communities, this suggests that deterministic selection is at work in similar environments. Consistent with our hypothesis, QPEN suggested that ecological drift was the dominant process for 75% of all pairwise comparisons among the final enrichment communities, while homogenous selection was the dominant ecological process for just 5%. Similarly, iCAMP indicated that drift was the dominant process for the majority of phylogenetic bins in 70% of the pairwise comparisons, while dispersal limitation and homogenous selection were dominant in 8% and 5% of the comparisons respectively.

As an orthogonal approach to assess the potential contribution of deterministic forces to the assembly of the enrichment communities, we performed consensus clustering analysis on the final communities after collapsing the taxonomy abundance table to the family level. Consensus clustering is a method of estimating how many subclasses (in this case, types of community structures) are present in a dataset (23), and has previously been used to infer whether deterministic forces contributed to variation across different enrichment communities (24). Specifically, if the number of inferred community types is much less than the number of samples, this would suggest some degree of determinism (non-randomness) in the community assembly process. Inspection of the delta area plot (in which lower values represent lesser gains in cluster stability with increasing *k*), as well as the cumulative distribution functions and heatmaps of the consensus matrices for different *k*’s (numbers of clusters), suggested that the optimal number of clusters was 9 (Fig. 3A-C, Fig. S2). However, clusters 5, 6, 7, and 9 were represented by only a single enrichment each. Organizing the taxonomy abundance bar plots by cluster identity suggested that cluster 1 (the largest cluster) was driven by a relatively high abundance of Methylocystaceae, whereas cluster 2 (the second-largest cluster) was driven by a high abundance of Comamonadaceae paired with a lower abundance of Methylocystaceae (Fig. 3D). Cluster 3 was appeared to be driven by a high abundance of ASVs that could not be classified beyond the domain Bacteria, cluster 4 was driven by a high abundance of Methylococcaceae, and cluster 8 was driven by a high abundance of Chromobacteriaceae. The samples in the four other clusters did not fall into any of these categories.

**Figure 3.**
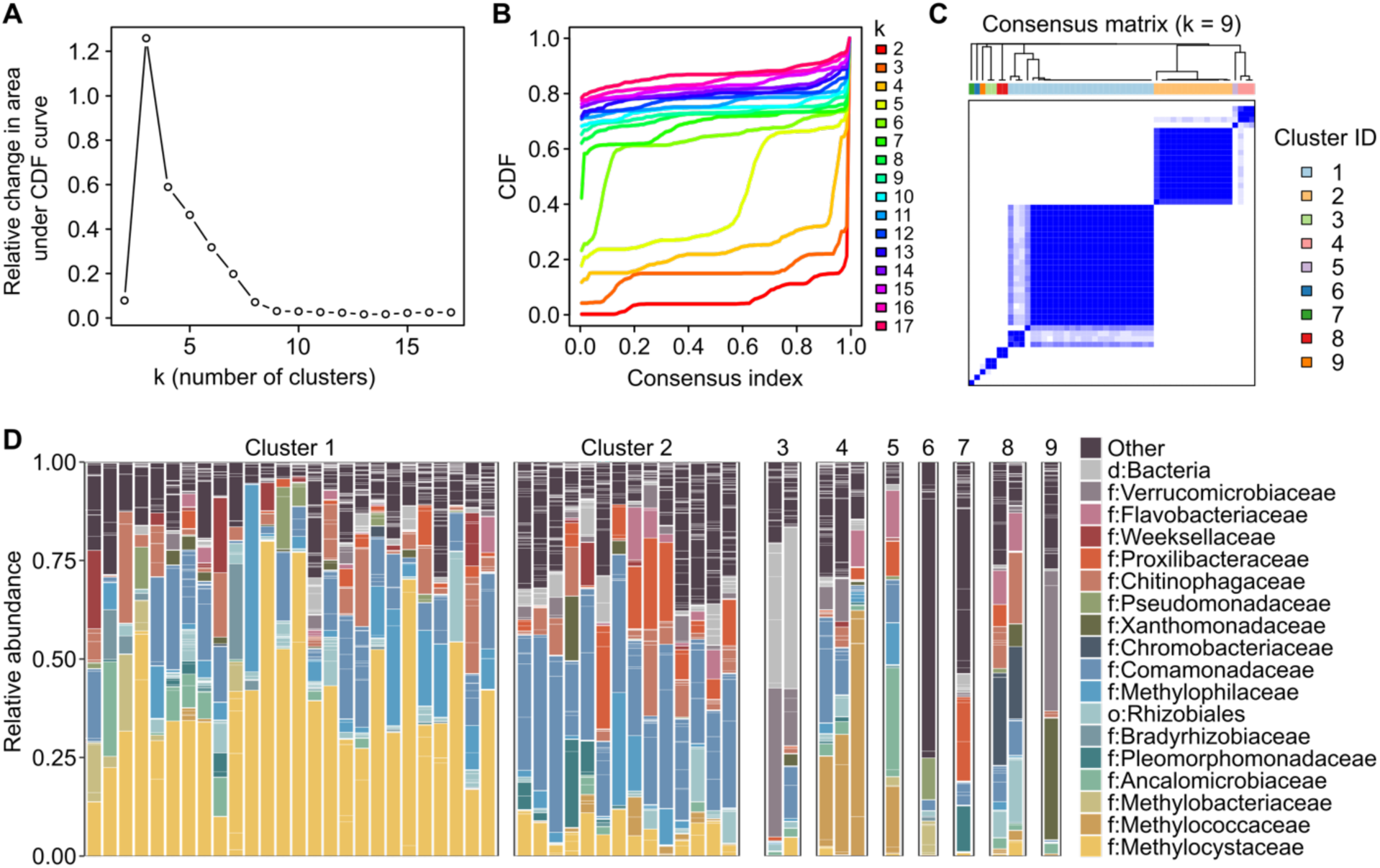
Consensus clustering analysis of enrichment communities. **A)** Delta area plot showing the relative change in area under the cumulative distribution function (CDF) curve for *k* versus *k* − 1. The optimal number of clusters is usually considered to be the value of *k* at which there is no longer an appreciable increase in area under the CDF. **B)** The CDFs of the consensus matrix for each *k* (indicated by colors), estimated by a histogram of 100 bins. **C)** The consensus matrix for *k* = 9. Darker blues in the matrix indicate higher consensus. **D)** Taxa bar plots for the samples in each consensus cluster for *k* = 9.

Taken together, these results suggest that despite the uniform culture conditions and theoretically strong selective force imposed by the use of minimal media with methane as the only added electron donor, stochastic processes (such as chance colonization—i.e. which strains were present at time zero—and random extinction) were important drivers of community assembly in our enrichments. Accordingly, even though certain types of community structures were more prevalent than others among the final enrichments (as revealed by the consensus clustering analysis), unpredictable and unique community structures also emerged.

### Putative parasitic bacteria belonging to the Candidate Phylum Radiation transiently bloomed in multiple independent enrichments

We were intrigued by the observation that ASVs that could not be classified beyond the domain Bacteria were abundant in several of the enrichments. This was particularly striking in the case of the final timepoint of the Staten Island root 2 10^−2^ enrichment, where the relative abundance of a single unclassified bacterial sequence, ASV607, was 40.9%. Besides ASV607, five other unclassified bacterial ASVs appeared at a relative abundance of >5% at least three times in our enrichment time course dataset: ASV611, ASV616, ASV684, ASV697, and ASV891 (Fig. 4A). To gain insight into the possible nature of these taxa, we used BLAST to search the respective ASV sequences against the NCBI core nucleotide database (Table S3). For ASV697, the best named hit was uncultured member of Chlamydiota detected in Chinese rice paddy soil, while for ASV684, the best named hit was an uncultured member of Lentisphaerota detected in lake sediment (also located in China), another environmental source of methane. For ASV616, there was no named hit with coverage >50%, but the top overall hits were from environments such as subsurface water, a sewage stabilization pond, and floodplain soil. For all three of the remaining sequences, the top named hits belonged to the members of the Candidate Phylum Radiation (CPR)—specifically, the candidate phyla Altimarinota (a.k.a. Gracilibacteria; ASV607 and ASV611) and Peregrinibacteriota (ASV891). Notably, the top overall hit for ASV891 was a 100% match to an unclassified 16S sequence from water that had been used to rinse rice grains; the second-best overall hit for ASV607 was a 97.83% match to an unclassified 16S sequence from the “heavy” isotopic fraction of activated sludge that had been incubated with ^13^C-labeled methane; and the top overall hits for ASV611 included sequences from lake sediment and a wetland. Taken together, these results suggest that bacteria closely related to the unclassified sequences in our enrichments might commonly occur in environments where methane is abundant.

**Figure 4.**
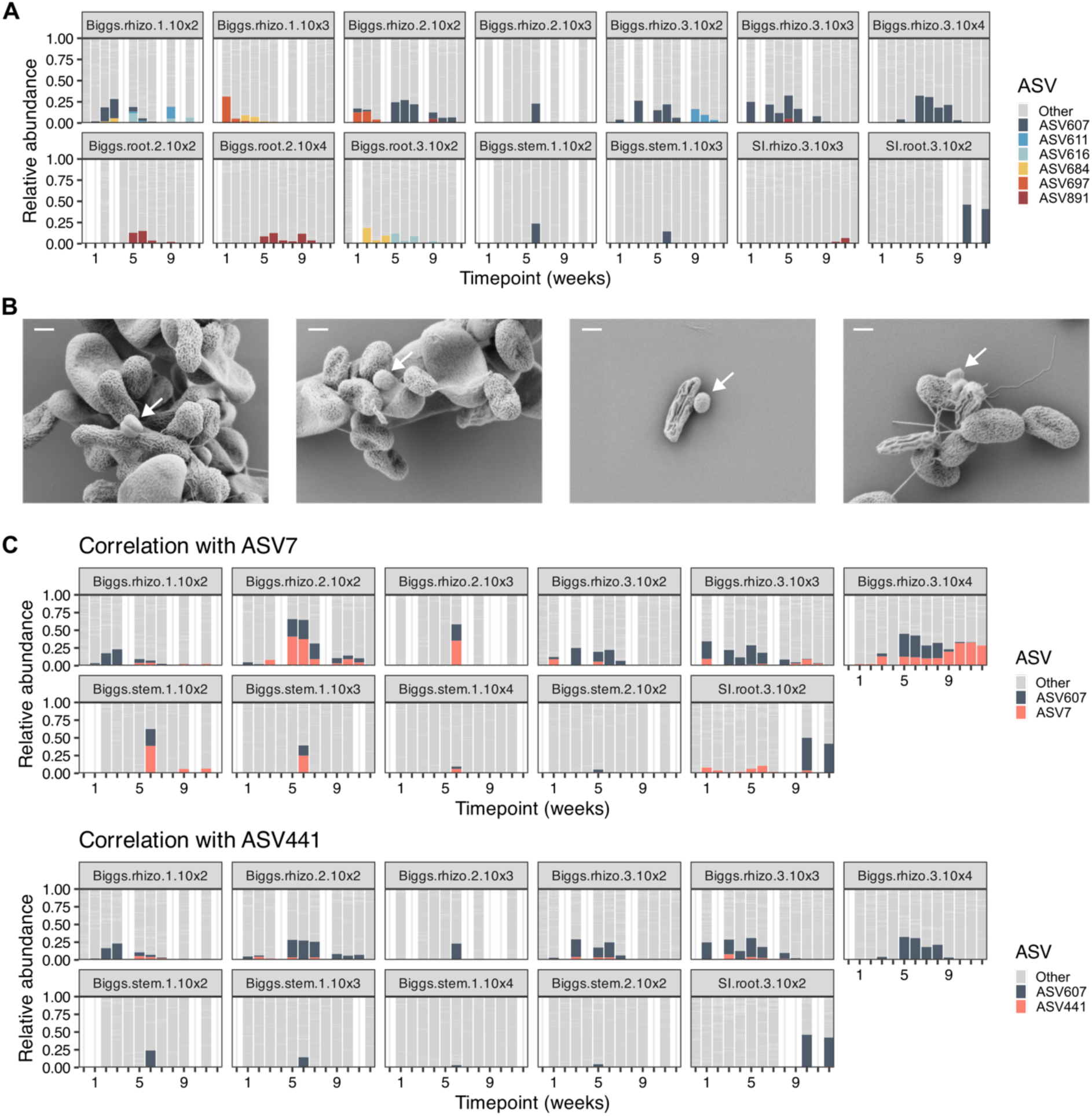
Presence of CPR bacteria in methane-fed enrichments. **A)** Taxa bar plot highlighting the six unclassified bacterial ASVs that appeared at a relative abundance of >5% at least three times (whether in different enrichments or multiple timepoints of a single enrichment). **B)** SEM images showing the putative CPR epibiont, marked with a white arrow. Scale bar = 300 nm. **C)** Taxa bar plots highlighting ASV607 along with the two candidates for its host based on correlation analysis.

Given that ASV607 was the most prevalent and abundant unclassified ASV in our enrichments (Fig. 4A), we focused on this taxon for further analysis. Members of the CPR, to which ASV607 putatively belongs, are mostly uncultivated but are thought to primarily be ultra-small (~200-500 nm diameter) episymbiotic parasites of other bacteria (25, 26). To determine whether such bacteria were present in a sample containing ASV607, we performed scanning electron microscopy (SEM) on the Staten Island root 2 10^−2^ enrichment after regrowing it from a frozen glycerol stock under the same conditions as the original enrichment; we focused on this enrichment because it had the highest relative abundance of ASV607 at the end of the original time course. Strikingly, we found multiple examples of such a bacterium in this culture: small (~340 nm diameter) cocci or diplococci that in all cases (*n* = 4) appeared to be directly attached to a larger host (Fig. 4B). Although we cannot be sure that these bacteria represent ASV607, no other 16S rRNA gene sequences belonging to the CPR or to other putative episymbionts were detected in the original enrichment. Together, these observations lend credence to the possibility that ASV607 is an episymbiont, similar to other studied members of the CPR.

Finally, we attempted to identify of the potential host(s) of ASV607 by looking for other ASVs whose abundances were correlated with this taxon. Since ASV607 might by chance not occur in every enrichment where its host was present, we first subsetted our enrichment time course dataset to only the enrichments where ASV607 was detected at least once. We then used two complementary methods of correlation analysis: SparCC, which was designed to handle some of the challenging characteristics of microbiome data (e.g. compositionality and sparsity) (27), and Pearson correlation, the method with the highest sensitivity for detecting parasitic interactions in a prior study based on simulated data (28). Importantly, the latter study also demonstrated that parasitic interactions between two taxa usually present as positive correlations (i.e. co-occurrence) rather than negative correlations in microbial abundance data (28). Only two ASVs were significantly positively correlated with ASV607 according to both methods: ASV7 (classified as Methylophilaceae) and ASV441 (classified as Spartobacteria). When we highlighted the abundances of these ASVs in the enrichments where ASV607 occurred, it became clear that ASV607 occurred at relatively high abundances in multiple enrichments where ASV441 was consistently below detection (Fig. 4C). By contrast, ASV7 was detected at least once in all of the enrichments where ASV607 was abundant, and at many timepoints these two taxa co-occurred with similar relative abundances (Fig. 4C). Thus, although we currently cannot rule out the possibility that ASV441 (or other taxa) could be a host for ASV607, ASV7 (Methylophilaceae) appears to be the most likely candidate for the primary host.

### Strains from the enrichments possess variable repertoires for metabolism of carbon and nitrogen

Given that the enrichment cultures were fed only with methane, the abundance and diversity of non-methanotrophic strains raised the question of what carbon sources could be supporting their growth. Methanotrophic activity has been shown to contribute to the pool of organic acids in rice paddy soils (12). Additionally, it has been previously reported that some methanotrophs can cross-feed methanol (the first product of methane oxidation) to non-methanotrophic methylotrophs (29). We therefore investigated whether organic acids and/or methanol could plausibly support the growth of members of the enrichment communities.

To do so, we isolated a total of 64 strains from the enrichments (Table S4). Of these isolates, three are methanotrophs (*Methylosinus* DMC42, *Methylosinus* DMC45, and *Methylocystis* DMC101), while the rest are non-methane-utilizing heterotrophs. The isolates represent 12 of the 16 families that had reached a final relative abundance of >10% in at least two enrichments (missing representatives only for Verrucomicrobiaceae, Methylococcaceae, Bradyrhizobiaceae, and Prolixibacteraceae). Three isolates belonging to the genus *Methyloraptor*, in the family Ancalomicrobiaceae, matched a similarly prevalent and abundant ASV that had been classified only to order level as Rhizobiales. We also obtained isolates for several other families that were less prevalent and/or abundant in the final enrichments, including Moraxellaceae, Sphingomonadaceae, Xanthobacteraceae, Caulobacteraceae, Devosiaceae, Microbacteriaceae, Sphaerotilaceae, and additional members of Rhizobiaceae (one of which had dominated a Biggs stem enrichment).

To obtain strain-level insights into the potential catabolic capabilities of the enrichment community members, we generated draft genome assemblies for a diverse selection of isolates representing most of the families in our collection. We then assessed the non-methanotrophic isolates’ genomic potential for the consumption of 62 common carbon sources, using the program GapMind (30). We considered a strain likely to be able to use a carbon source if GapMind identified a complete pathway with only high-confidence hits, and a “maybe” if GapMind identified a complete pathway but at least one step was at best a medium-confidence hit. Strains were considered unlikely to be able to use a carbon source if any of the best hits in the respective pathway were low-confidence, i.e. “gaps.” This analysis revealed that nearly all of the sequenced isolates are likely to be able to use fumarate as a carbon source, and most of the isolates (with the exception of members of Methylophilaceae and *Cloacibacterium*) are also likely to be able to use multiple other organic acids, such as succinate, malate, and acetate (Fig. 5; Table S5). By comparison, the capacity to utilize sugar-related compounds and amino acids as carbon sources appeared to be less widespread.

**Figure 5.**
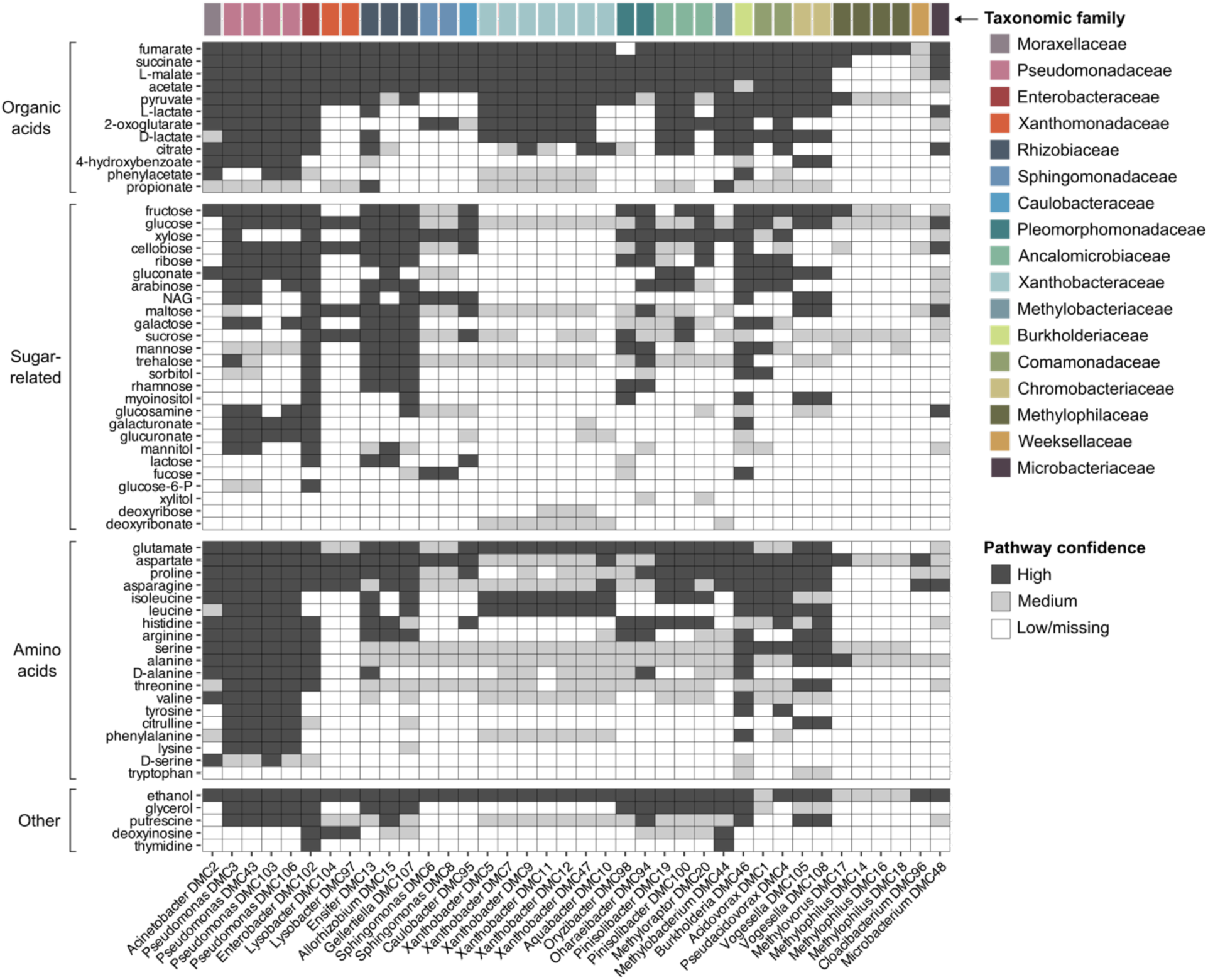
Putative carbon source utilization pathways in non-methanotrophic isolates from methane-fed enrichments. The presence of pathways for utilizing 62 common carbon sources was assessed for 38 isolate genomes using GapMind. Pathways were considered high, medium, or low confidence/missing according to the lowest confidence level for any step in the pathway (e.g. a pathway in which only a low-confidence hit was found for at least one step was considered low confidence overall).

We also assessed whether the non-methanotrophic isolates could potentially utilize methanol, which is not included in GapMind. To do so, we searched the genomes for homologs of the four known types of methanol dehydrogenases (calcium-dependent Mxa; lanthanide-dependent Xox; the type I alcohol dehydrogenase Mdh2; and NAD-dependent Mdh) (Fig. 6A; Table S5). For strains that possessed at least one putative methanol dehydrogenase, we also looked for homologs of the genes needed for the ribulose monophosphate (RuMP) cycle, the serine cycle, and the Calvin cycle, which are the three known pathways for assimilation of C1 units by methylotrophic bacteria (31) (Fig. 6B; Table S5). The results were mostly consistent with what has been reported for the respective taxonomic groups: members of *Methylovorus*, *Methylophilus*, *Methylobacterium*, *Methyloraptor, Oharaeibacter*, and *Xanthobacter* are all known methylotrophs, and all sequenced isolates from these genera possessed both Mxa and Xox, as well as a complete C1 assimilation pathway. However, one isolate from a genus not known to be methylotrophic also showed possible genomic capacity for methylotrophy: *Pinisolibacter* sp. DMC19 possesses a homolog of XoxF (though it seemingly lacks other Xox genes), as well as a complete Calvin cycle. Another *Pinisolibacter* isolate also possesses XoxF and a near-complete Calvin cycle, but appears to be missing glyceraldehyde-3-phosphate dehydrogenase; while *Oryzibacter oryziterrae* DMC98 possesses XoxF along with a near-complete serine cycle, but appears to be missing phosphoenolpyruvate carboxylase. A few other isolates from genera not typically known to consume methanol contained putative homologs of XoxF, Mdh2, and/or NAD-dependent Mdh, but lacked multiple steps in each of the C1 assimilation pathways, suggesting that they are unlikely to use methanol as a carbon source.

**Figure 6.**
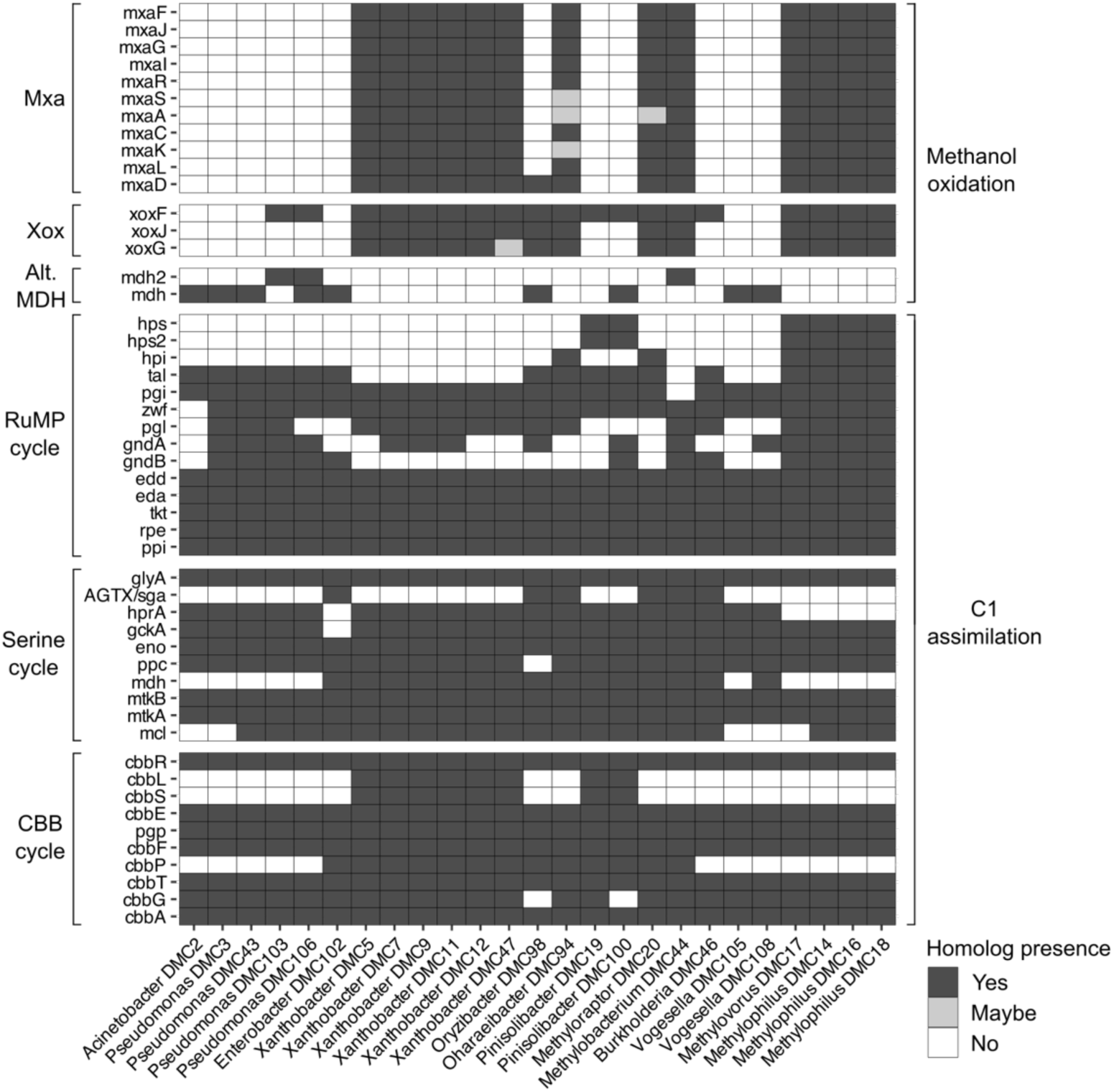
Genomic capacity for methanol catabolism in non-methanotrophic isolates from methane-fed enrichments. Presence of homologs of known methanol dehydrogenases and the genes needed for the three known bacterial C1 assimilation pathways. “Alt. MDH” = alternative methanol dehydrogenase other than the classical Mxa and Xox types. Only strains possessing at least one putative methanol dehydrogenase are shown. Homologs were detected using DIAMOND for methanol dehydrogenases (cutoffs: >40% identity and >80% coverage, with “maybe” denoting a hit with 37-39% ID and >80% coverage, or >40% identity and 75-79% coverage), and HMMER for C1 assimilation genes (cutoffs: e-value < 10^−5^ and coverage >80%).

Finally, given that methanotrophic activity has previously been shown to impact nitrogen cycling in rice paddy soils (12, 17), we assessed the genomic capacity for each isolate, including methanotrophs, to fix nitrogen and/or perform denitrification. Several strains possessed *nifD*, *nifK*, and *nifH*, indicating that they can likely fix nitrogen (Fig. 7; Table S5); these belonged to the genera *Pseudacidovorax*, *Burkholderia*, *Xanthobacter*, *Aquabacter*, *Pinisolibacter*, *Methyloraptor*, *Methylophilus*, and *Methylosinus* (which is methanotrophic). Among these, to our knowledge the nitrogen-fixing potential of *Pinisolibacter* and *Methylophilus* has not been previously reported (although a metagenome-assembled genome classified under the family Methylophilaceae, to which *Methylophilus* belongs, was recently found to contain a *nif* locus (32)). Only two out of three sequenced isolates of *Methylophilus* possessed the *nif* genes, indicating that nitrogen fixing capabilities vary within this genus. As for denitrification potential, we found a variety of combinations of genes involved in the different steps of denitrification (Fig. 7; Table S5). Only *Pinisolibacter* sp. DMC100 possessed a complete denitrification pathway. Some strains appeared to be capable of only nitrate reduction; others only had the genes for nitrous oxide reduction. Still others lacked the genes for nitrate reduction but possessed all other steps of denitrification, or conversely had all steps except for nitrous oxide reduction. Additionally, three strains only possessed the genes for non-consecutive steps (nitrate and nitric oxide reduction, nitrate and nitrous oxide reduction, or nitrite and nitrous oxide reduction).

**Figure 7.**
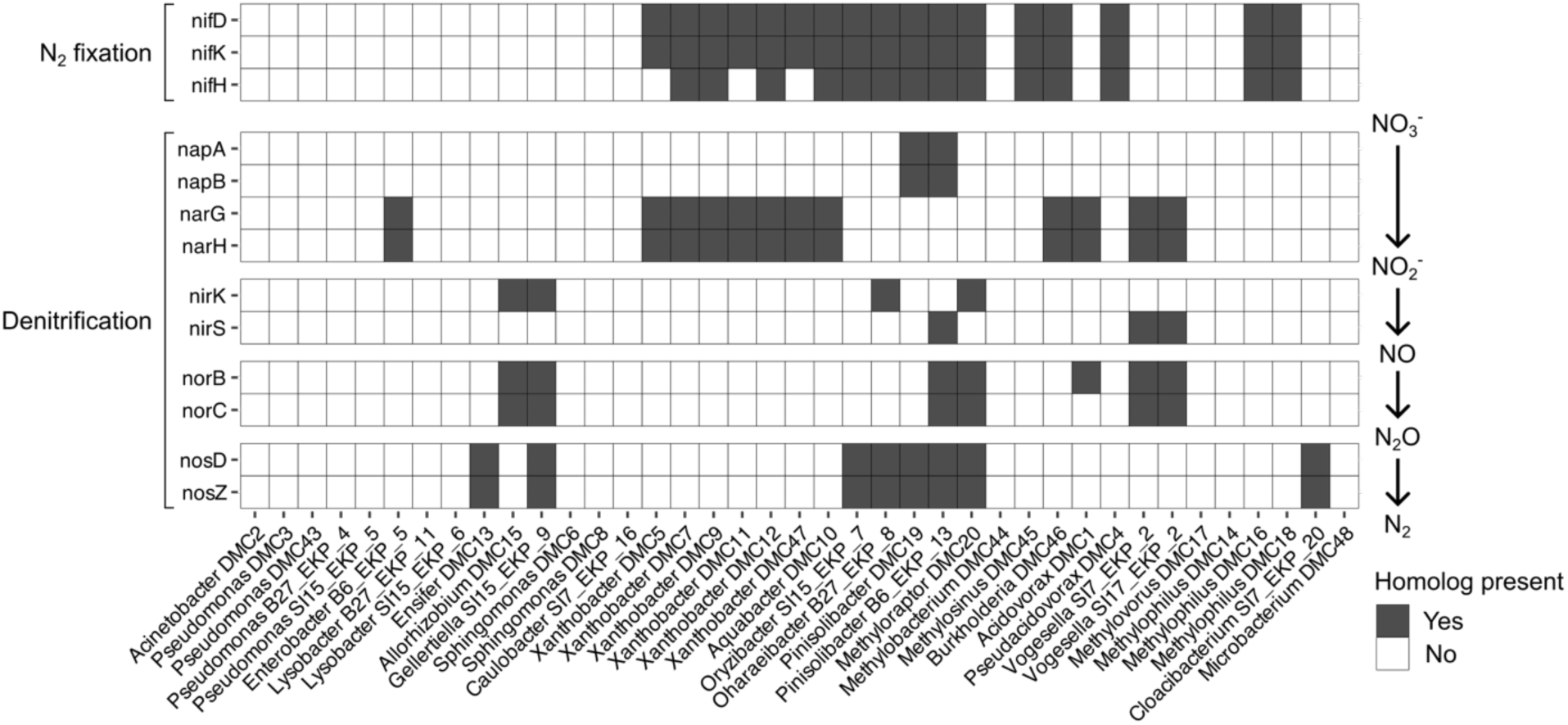
Genomic capacity for nitrogen fixation and denitrification in isolates from methane-fed enrichments. Presence of homologs of key genes needed for nitrogen fixation and the four steps of denitrification. Homologs were detected using microTrait on KBase. Nap and Nar are alternative nitrate reductases—i.e. only one of the two is needed to reduce nitrate to nitrite. Similarly, for reduction of nitrite to nitric oxide, either NirK or NirS is sufficient.

## Discussion

In this study, we utilized methane-fed enrichment cultures to determine which bacterial members of the rice microbiota can be supported by methane-derived carbon, revealing considerable taxonomic diversity among the potential beneficiaries of methanotrophic activity. We found that the composition of rice-derived, methane-supported microbial communities is significantly affected by the type of plant part used as the inoculum (with stem enrichments exhibiting lower alpha diversity than root and rhizosphere enrichments), but is less affected by field site (Fig. 2, Table 1). The lower alpha diversity of the stem enrichments is consistent with prior studies indicating that the stem or leaf microbiotas of rice and other plants tend to have lower alpha diversity compared to the rhizosphere and roots (33, 34). While many of our enrichments had a high abundance of type II methanotrophs and/or members of Comamonadaceae, a variety of overall community structures emerged (Fig. 1, Fig. 3), suggesting a limited contribution of deterministic forces to community assembly during growth on methane. Finally, genomic analysis indicated that organic acids and methanol are likely carbon sources that may support the growth of many non-methanotrophs in methane-fed communities (Fig. 5, Fig. 6). Taken together, these results shed new light on the contribution of methanotrophs to the rice microbiome.

The diversity of non-methanotrophic taxa that were enriched alongside methanotrophs in our cultures, representing at least 14 families, greatly expands the knowledge of which taxa may benefit from methanotrophic activity in rice paddies. In prior DNA-SIP studies where rice roots or paddy soils were incubated with isotopically-labeled methane, relatively few taxa were identified (10–12), possibly due to the limited sensitivity of this technique and the presence of alternative carbon sources in those experimental setups. In our enrichments, as in the DNA-SIP studies, some carbon from the soil or plant material would have been present at the start of the experiment, but any such nutrients would have then been diluted by the two sequential transfers, if not already fully consumed in the first few weeks of incubation. Thus, with the possible exception of a few enrichments where the final abundance of putative methanotrophs appeared to be extremely low (which may have occurred due to stochastic extinction), taxa that were abundant at the final timepoint of our enrichments were likely primarily supported by methane-derived carbon. Among the non-methanotrophic taxonomic groups that were particularly prevalent and/or abundant in our enrichments, Comamonadaceae, Methylophilaceae, Rhizobiales, and Sphingomonadales have also been labeled with methane-derived carbon in DNA-SIP studies (10, 12), providing further evidence that these taxa genuinely benefit from methanotrophic activity in rice fields.

The heterogeneity of our enrichments indicated a significant contribution of stochastic ecological forces to community assembly during growth under methane. While this may in part have been driven by the particulars of our experimental setup (e.g., the use of a dilution series, and the relatively long four-week incubation time between transfers), the variety of observed community structures, together with the diversity of the enriched taxa, suggests that the impact of altering methanotrophic activity in rice microbiomes might not be easily predicted. That being said, our enrichments also bore some taxonomic resemblance to previous studies on methane-supported microbial communities from other environments. For example, *Methylophilus*, *Pseudomonas*, and *Flavobacterium*, three prominent genera in our enrichments (Fig. 1), were also prevalent in methane-fed enrichments from lake sediment (35, 36). Moreover, a study on naturally-occurring methane-supported biofilms in a mineral spring cave reported the presence of Flavobacteriales, Verrucomicrobia, Rhizobiales, Sphingomonadales, Pseudomonadales, and Xanthomonodales, among other non-methanotrophic taxa (37), all of which were abundant in at least two of our enrichments (note that Verrucomicrobia also encompasses methanotrophic acidophiles belonging to the genera *Methylacidiphilum* and *Methylacidimicrobium* (38), but these were not detected in either the mineral spring cave or our enrichments). Thus, it appears that the assembly of methane-supported communities is not entirely random, and many of the taxonomic groups in our enrichments may be associated with methanotrophs not only in rice paddies, but also more broadly.

Another notable finding was the prevalence of CPR bacteria in our enrichments (Fig. 4, Table S3). To our knowledge, this phenomenon has not previously been explicitly reported in a methane-supported microbial ecosystem. However, many of the closest sequenced relatives for the putative CPR bacteria in our enrichments were also derived from environments where methane is abundant, indicating that our detection of CPR members in methanotrophic communities is not an anomaly. Given that the DNA of one such organism was labeled with methane-derived carbon in a DNA-SIP study (see Table S3), some of these bacteria may be genuine beneficiaries of the activity of methanotrophs. This interpretation is further supported by the positive correlation between ASV607 (a putative CPR member) and ASV7 (a member of the methylotrophic family *Methylophilaceae*) in our enrichments (Fig. 4C). Importantly, standard 16S primer pairs are a poor match for many CPR members (39). Thus, these organisms may be even more prevalent and abundant in rice microbiomes and other methane-supported microbial communities than is reflected in currently available data. Detailed studies on CPR bacteria are challenging due to the difficulty of isolating these organisms. Nevertheless, future attempts to confirm the identity of the host(s) for rice-associated CPR bacteria, and to ascertain the nature of the host-epibiont relationship (parasitic, predatory, or commensal), may help improve the understanding of rice microbiome dynamics.

Regarding the question of how non-methanotrophic growth is supported in methane-fed communities, the prevalence of pathways for organic acid catabolism in isolates from our enrichments (Fig. 5) is consistent with the idea that organic acids are likely a key class of cross-fed metabolites, corroborating prior evidence that methanotrophs contribute to organic acid production in paddy soils (12). Whether this contribution is primarily direct or indirect remains unclear, but certain type I and type II methanotrophs have been shown to excrete organic acids in pure cultures during growth on methane under low oxygen conditions (40–42). Low oxygen conditions likely prevailed in our enrichment cultures over time, as the initial molar ratio of oxygen to methane in the headspace was less than 1 (see Materials and Methods), whereas two oxygen molecules are required to oxidize one molecule of methane all the way to carbon dioxide (43). Additionally, the abundance of several methylotrophic taxa in our enrichments (Fig. 1, Fig. 6) suggests that methanol may also have been cross-fed from methanotrophs, as has been previously demonstrated in pairwise co-cultures of a methanotroph and a methylotroph (29). However, organic acids and methanol may not be the only cross-fed carbon sources in methanotroph-supported communities. Several taxa enriched in our cultures belong to the phylum Bacteroidota, members of which are best known for their ability to degrade complex carbohydrates (44). Consequently, it has previously been suggested that Bacteroidota members that frequently co-occur with methanotrophs (e.g., *Flavobacterium* species) may be supported by extracellular polysaccharides produced by other community members, though direct evidence of such interactions is still lacking (5). These taxa proved difficult to isolate from our enrichments, and thus we only have one representative genome sequence so far, from a strain of *Cloacibacterium*. Potentially consistent with the extracellular polysaccharide hypothesis, this strain does not possess high-confidence pathways for the usage of either organic acids or methanol (Fig. 5), suggesting that it may indeed have been supported by other compounds in the enrichment cultures. We also cannot rule out that nutrients may also have been released by cell lysis over the course of our experiment, potentially supporting some of the taxa that only persisted at a very low relative abundance in the enrichments. While beyond the scope of the current study, future work will be directed at more precisely identifying the carbon sources exchanged among members of methanotroph-supported microbial communities.

Finally, the presence of genes related to nitrogen cycling in the majority of isolates from the enrichments is consistent with previous indications of functional links between methanotrophy, nitrogen fixation, and denitrification in rice paddy microbial communities (12, 45). Whether the activity of methanotrophs tends to stimulate nitrogen fixation or denitrification more strongly *in situ* is unknown; the balance between the two presumably depends not only on the makeup of a given community, but also environmental parameters such as local oxygen gradients and application of fertilizers (45, 46). With respect to denitrification, we observed that many strains possessed only partial pathways (Fig. 7), as is common in many natural environments (47), suggesting that cross-feeding of not only carbon sources (electron donors) but also denitrification intermediates (electron acceptors) may occur in methanotroph-supported communities under low oxygen tensions. As mentioned above, oxygen tension was likely low in our enrichments for much of the time course, making nitrate an important alternative electron acceptor that could have shaped the emergent community structures.

In summary, our findings suggest that methanotrophs can help support a wide range of metabolically-diverse heterotrophic bacteria in the microbial ecosystem of rice paddies, with the potential for impacts on important ecosystem functions such as nitrogen fixation and denitrification. Furthermore, our collection of 64 taxonomically-diverse isolates, more than half of which have draft genomes, represents a valuable resource for future mechanistic studies on how rice-associated methanotrophs interact with their native neighbors. Given that methanotrophs affect both carbon cycling and nitrogen cycling in rice paddies, it will be worthwhile to continue gaining a deeper understanding of how methanotrophic bacteria influence the structure and function of the rice microbiome under different conditions, so as to support the long-term sustainability and productivity of this globally-important staple crop.

## Acknowledgements

This work was supported by the Laboratory Directed Research & Development program at Lawrence Berkeley National Laboratory [award number 23-071 to D.C.]. We thank the Sundaresan and Schuppenhauer labs for providing the field-grown rice plants, Deepika Awasthi for assistance with establishing a methane gassing station, Morgan Price for assistance with data processing, and Misun Kang and the staff at the University of California Berkeley Electron Microscope Laboratory for advice and assistance with electron microscopy sample preparation and imaging.

## Materials and Methods

### Time course of methane-fed enrichment cultures

Six rice plants were collected from two fields (three plants per field), located in Biggs, California (39.4643, −121.7340) and Staten Island, California (38.1235, −121.5490), in the summer of 2022. The Biggs samples were collected in late July, and the Staten Island samples were collected in late August. Each plant was collected from a different area of the respective field and had the entire root structure dug up with the attached soil. The collected plants were placed in individual clean zipper-top plastic bags in 5-gallon buckets and transported back to the laboratory. The protocol for establishing methane-fed cultures from the rhizosphere was based on previous work (48), with modifications made to include the root and stem. For each plant, after removing soil not tightly bound to roots, a portion of the roots was dissected and placed in a 15 mL conical tube containing 10 mL of dilute nitrate mineral salts (dNMS) media, made according to the recipe of (48) except that the vitamin solution was procured from the American Type Culture Collection (catalog no. MD-VS). The complete media composition is provided in Table S6. The roots were vigorously vortexed to generate the rhizosphere samples. Then, a portion of the roots was ground in 1 mL dNMS for 45 s using a sterilized stainless-steel grinding jar (Qiagen, catalog no. 69985) in a Qiagen TissueLyser II. Stem samples were processed in a similar manner. 0.5 mL of the homogenized samples, representing the week 0 time point, was spun down at 4000 rpm for 20 min in a tabletop centrifuge, and the pellets were stored at −20°C until DNA extraction. The remainder of each sample was diluted 10^−2^, 10^−3^, and 10^−4^ in dNMS (Staten Island stem samples were only diluted up to 10^−3^), and 0.5 mL of the resulting suspensions was injected into sealed, autoclaved Balch tubes containing 5 mL of dNMS. Approximately 5 mL of the headspace was removed by syringe and replaced with 5 mL of methane to create a 20% methane headspace. The cultures were incubated upright with shaking at 24°C. Every seven days, 0.5 mL was collected from each culture and pelleted for later DNA extraction as above. Additionally, 70 µL of each culture was used to make a frozen stock in 15% glycerol, stored at −80°C. At the same time, 5 mL of headspace was withdrawn and replaced with another 5 mL of methane. Every four weeks, 0.5 mL of each culture was transferred to a new Balch tube containing 5 mL of fresh dNMS. The experiment was terminated after a total of 12 weeks. Note that in this setup, the cultures were most likely electron acceptor-limited rather than methane-limited. The initial quantities of methane and oxygen per tube were 210 µmol and 140 µmol respectively, whereas two moles of oxygen are required to completely oxidize one mole of methane to carbon dioxide (43), involving the transfer of 8 electron equivalents (dioxygen can accept 4 electron equivalents per mole). Even if nitrate was used as an alternative electron acceptor by some community members, only 12.5 µmol of nitrate was supplied (which can accept 5 electron equivalents per mole via complete denitrification).

### 16S rRNA gene amplicon sequencing

Genomic DNA was extracted from pelleted samples using the Qiagen DNeasy Blood & Tissue kit (catalog no. 69506), following the manufacturer’s protocol for Gram-positive bacteria. The extracted DNA was PCR amplified using dual-index 16S V4-V5 primers (515F/926R) (49). A peptide nucleic acid (PNA) PCR clamp was used to block the amplification of plastid and mitochondrial DNA (50). The resulting libraries were quantified using the Qubit dsDNA BR Assay kit (Invitrogen, catalog no. Q32853), pooled, cleaned using a PCR purification kit, and sequenced on an Illumina MiSeq (PE 2x300). The paired-end reads were assembled using PEAR (51), and reads with more than one expected error were detected using USEARCH (52) and discarded, along with rare sequences (fewer than 4 reads in any sample). Chimera removal and error correction were performed with the UNOISE3 algorithm (53). Taxonomy was assigned using SINTAX (54) with the RDP training set v18.

### 16S sequencing data analysis

The following initial analysis steps were performed in Qiime2 (version amplicon-2024.5): Samples were rarefied to 900 reads (based on the histogram of reads per sample), a phylogenetic tree was constructed using MAFFT alignment and FastTree, core alpha and beta diversity metrics were calculated, and principal coordinate analysis (PCoA) matrices were generated. SparCC and Pearson correlation analyses were also performed in Qiime2, using the SCNIC plugin (55). All other analyses were performed in R. PERMANOVA analyses were performed with the adonis2 function from the package vegan (version 2.6-10) (56). PERMDISP tests were performed with the permutest function in conjunction with the betadisper function, also from vegan. When testing the effect of time point, permutations were constrained within each enrichment and the data were treated as a series. When testing the effect of field site or sample type, permutations were constrained within time point. Assessment of the contribution of stochastic versus deterministic processes during community assembly was performed using the functions qpen and icamp.big in the iCAMP package (20). Consensus clustering was performed using the package ConsensusClusterPlus, with maxK = 17, reps = 100, pItem = 0.8, pFeature = 1, and clusterAlg = “hc” (57). Except for the consensus clustering plots, which were produced by the ConsensusClusterPlus package, all plots were made with the R package ggplot2 (58).

### Scanning electron microscopy

To prepare the scanning electron microscopy (SEM) sample (Electron Microscopy Laboratory, University of California, Berkeley), the Staten Island root 2 10^−2^ enrichment was regrown, from a frozen glycerol stock, in dNMS with 20% methane (shaking at 125 rpm at 25°C). The regrown culture was sub-cultured twice at a 1:100 dilution ratio on day 15 and day 23. On day 29, 10 mL of the culture was pelleted (5000 g for 10 min), and the cells were gently resuspended in a fixative solution (2% glutaraldehyde in 0.1M sodium cacodylate buffer, pH 7.2). The sample was placed at 4°C on a rotator overnight and subsequently stored at 4°C in the dark (standing) until further processing. Prior to imaging, the sample was washed 3x in 0.1M sodium cacodylate buffer, pH 7.2, postfixed 3x in 1% osmium tetroxide in 0.1M sodium cacodylate buffer, pH 7.2 for 1 h, and then washed 3x in 0.1M sodium cacodylate buffer, pH 7.2. Samples were dehydrated in a graded ethanol series up to 100%. Samples were then transferred onto 0.1% (w/v) poly-L-lysine– coated coverslips and subjected to critical point drying (Tousimis, Mayland, USA). The dried coverslips were mounted onto aluminum stubs using conductive carbon tape and sputter-coated with a gold/palladium alloy (Safematic GmbH, Switzerland). SEM imaging was performed using a Zeiss Crossbeam 550 (Carl Zeiss Microsystems GmbH, Oberkochen, Germany).

### Isolation of strains from the enrichment cultures

Strains were isolated from the original enrichment cultures, as well as enrichments regrown from frozen glycerol stocks, by plating a 10-fold dilution series onto dNMS with 1.5% agarose and 0.2% methanol; dNMS agarose under a mixed methane/air atmosphere; or 0.1x R2A with 1.5% agar. To obtain the *Methylocystis* isolate, a 10-fold dilution series (10^−1^ to 10^−8^) of an enrichment regrown from the frozen stock was set up in a 48-well plate and incubated statically under 20% methane at room temperature for 4 weeks; then, a loopful of the culture in the 10^−8^ dilution well was streaked onto dNMS agarose and incubated under a mixed methane/air atmosphere. The mixed methane/air atmosphere was achieved by flooding a mostly-closed BD EZ Incubator Container with methane for 30-60s and then quickly latching the lid. All plates were incubated at room temperature, and morphologically distinct colonies were picked and restreaked 3-4 times until the colony morphologies appeared uniform. The 16S rRNA gene was then amplified by colony PCR, using Platinum Hot Start Master Mix (Invitrogen) with the primers 27F (AGAGTTTGATCMTGGCTCAG) and 1492R (TACGGYTACCTTGTTACGACTT). The PCR products were Sanger sequenced at Elim Biopharm, Inc. with the same 27F and 1492R primers. The resulting forward and reverse sequences were aligned in Geneious Prime and subjected to BLAST against the NCBI core nucleotide database.

### Genomic analysis of isolated strains

Genomic DNA was isolated from selected strains using the Qiagen DNeasy Blood & Tissue kit (catalog no. 69506), following the manufacturer’s protocol for Gram-positive bacteria. Library preparation and whole-genome sequencing was performed by Novogene on an Illumina NovaSeq X Plus. The raw reads were cleaned with Trimmomatic on KBase (59), using the default settings. Draft genomes were assembled with SPAdes (v3.15.3), with the minimum contig size set to 1000 bp, and annotated with Prokka (v1.145), also on KBase. Genomes were checked for completeness and contamination with CheckM. Taxonomy was assigned using the Microbial Genomes Atlas pipeline against the Genome Taxonomy Database (GTDB, version r214) (60, 61). Annotation of common carbon source utilization pathways was performed using the online GapMind tool (30). Detection of homologs of known methanol dehydrogenases was performed using DIAMOND with the “sensitive” setting (62); a hit was considered a homolog if it had >40% identity to the reference sequence, with at least 80% coverage. The list of reference methanol dehydrogenase sequences was obtained from a prior publication (63). For C1 assimilation pathways, DIAMOND with the same settings failed to detect complete pathways in some of the known methylotrophs. We therefore instead used HMMER (version 3.4) (64) with the HMM profiles from the same prior publication (63), requiring hits to have at least 80% coverage of the respective model to be considered a homolog. Genes related to nitrogen cycling were detected using the application “Search with HMMs of MicroTrait Bioelement families - v1” on KBase (59, 65).

### Data availability

The raw sequence data for 16S rRNA gene amplicons from the enrichments are available on the NCBI Sequence Read Archive (SRA) under the accession PRJNA1310742. Draft genome assemblies have been deposited DDBJ/ENA/GenBank under the accession PRJNA1310743.

