## Supplemental Figures for "Methane-fed microbial communities enriched from field-grown rice support diverse heterotrophic bacteria"

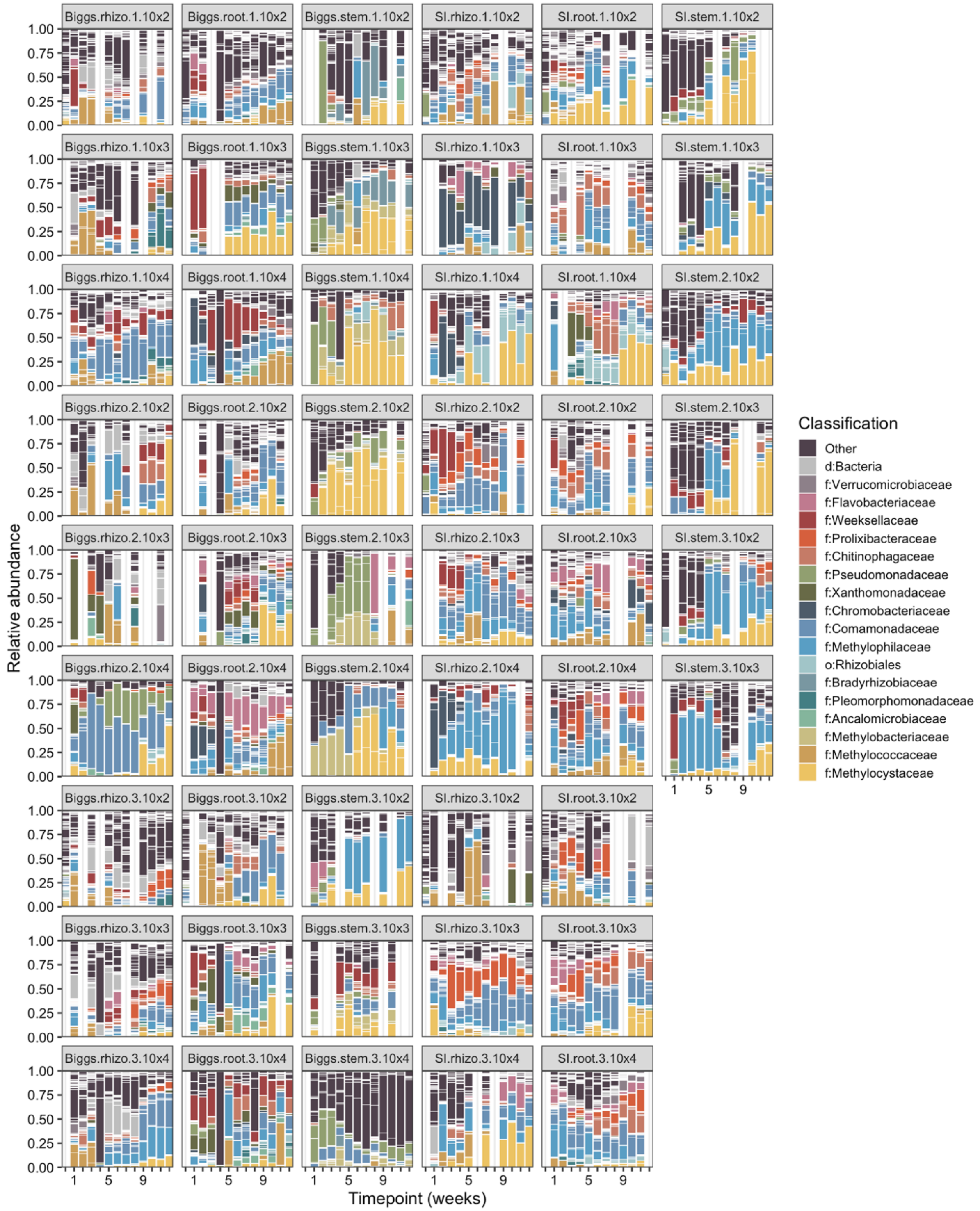


**Supplementary Figure 1.** Taxa bar plots for the weekly samples from all 51 enrichment cultures over the 12-week time course. Sample naming scheme: [field of origin, where SI = Staten Island].[plant part].[plant replicate number].[inoculum dilution factor, where 10x2, 10x3, and 10x4 are shorthand for 10^-2, 10^-3, and 10^-4]. Empty columns reflect insufficient reads for some of the samples. In the legend, d = Domain, o = Order, f = Family.

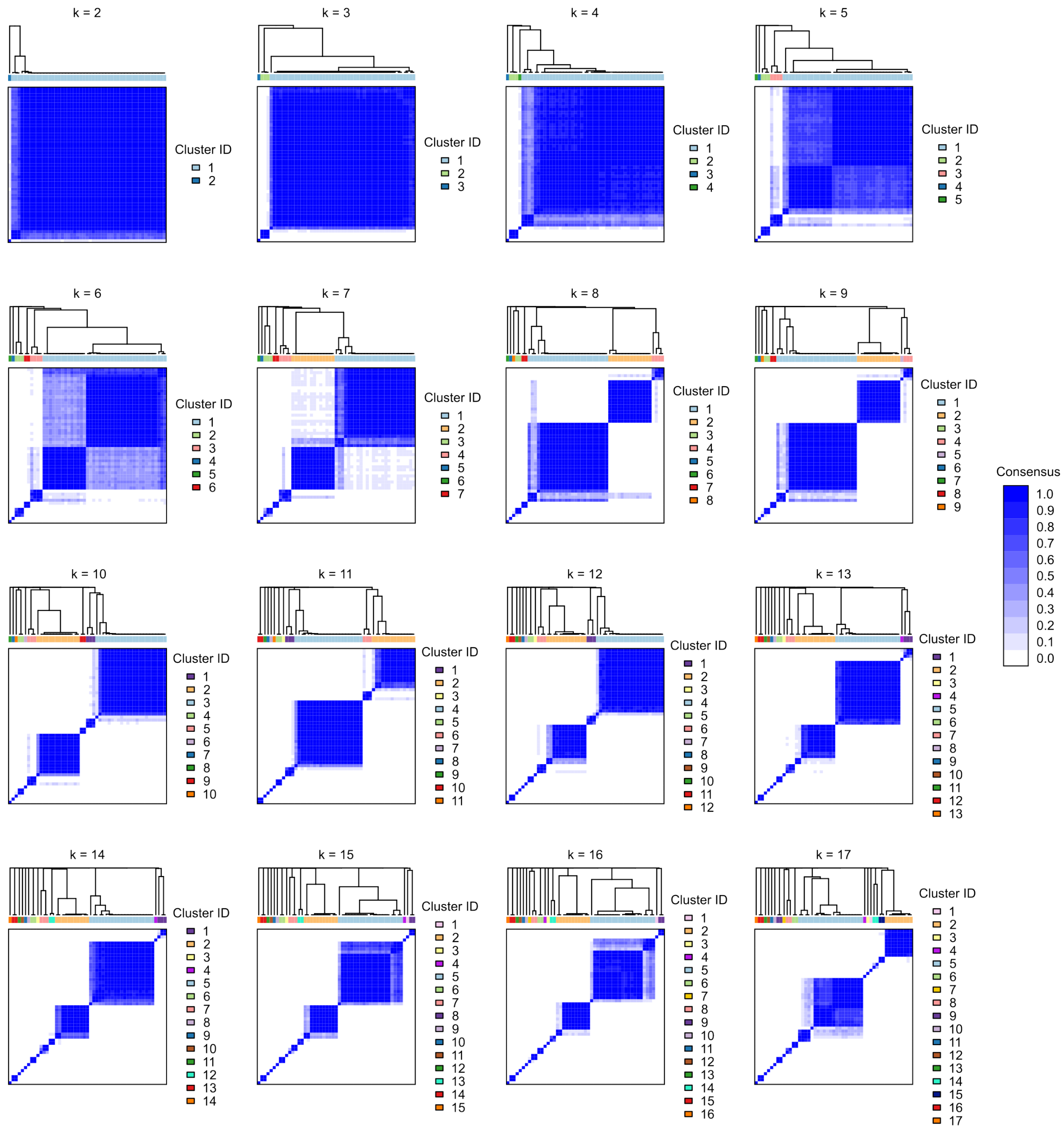


**Supplementary Figure 2.** Consensus matrices for *k* = 2 through *k* = 17, based on the family-level taxonomy of the enrichment communities at the final timepoint.
